# Preliminary Evidence for Anxiety-Linked Neural Sensitivity to Emotional Faces Using Fast Periodic Visual Stimulation

**DOI:** 10.1101/2025.02.15.638321

**Authors:** David Vandenheever, Haleigh Davidson, Jennifer Kemp, Zack Murphy, Autumn Kujawa, Jingyi Shi, Michael R. Nadorff, Kayla Bates-Brantley, MacKenzie Sidwell

**Affiliations:** Neural Engineering Research Division (NERD), Mississippi State University, Mississippi; Department of Psychology and Human Development, Vanderbilt University, Nashville, Tennessee; Department of Mathematics and Statistics, Mississippi State University, Mississippi; Department of Psychology, Mississippi State University, Mississippi; Department of Counseling, Higher Education Leadership, Educational Psychology and Foundation, Mississippi State University, Mississippi

**Keywords:** EEG, FPVS, Anxiety, internalizing disorder, ERP

## Abstract

Facial expression processing is crucial for social communication and survival, with anxiety disorders often linked to alterations in attentional biases toward threat-related stimuli. While previous studies using event-related potentials (ERPs) have yielded conflicting findings regarding threat sensitivity in anxiety, Fast Periodic Visual Stimulation (FPVS) offers a high signal-to-noise, implicit alternative for assessing emotion processing. This study utilized FPVS to investigate neural responses to facial expressions in individuals with elevated anxiety-related characteristics (e.g., prior diagnosis or elevated symptom scores) and those without. EEG responses were recorded while participants viewed sequences of neutral faces interspersed with emotional oddball expressions (angry, fearful, happy, and sad). Results revealed robust neural discrimination responses to all facial expressions. Participants with anxiety-related characteristics showed significantly greater summed baseline-corrected amplitudes (BCA) at occipital electrodes in response to angry and sad oddball faces compared to the low-anxiety group. This was accompanied by reduced top-down interactions. Although, dimensional anxiety scores were generally low, suggesting results may reflect residual or trait-level differences rather than acute symptomatology, these findings provide preliminary evidence that FPVS may be sensitive to enduring differences in emotion processing associated with anxiety vulnerability.

## 1. Introduction

The rapid and accurate processing of facial expressions is fundamental to human social interaction, as it enables the detection of emotional cues that are critical for survival. From an evolutionary standpoint, the ability to swiftly identify and differentiate facial expressions, particularly those signaling potential threats such as anger or fear, is vital for adaptive behavior [1]. Humans have evolved complex neural systems to categorize facial expressions efficiently, facilitating both rapid visual discrimination and the generalization of emotional cues across different facial identities [2].

Individuals with anxiety disorders often exhibit heightened sensitivity to negative and threat-related stimuli, leading to an exaggerated attentional bias toward such cues. This bias is believed to play a significant role in the development and maintenance of anxiety disorders [3]. Supporting this, studies utilizing electroencephalography (EEG) and event-related potentials (ERPs) have consistently demonstrated enhanced neural responses to threatening and negative facial expression in anxious individuals [4]. Specifically, anxiety has been associated with increased amplitudes in early ERP components such as the P1, P2, N170, and P200, reflecting heightened early-stage processing of negative and threat-related stimuli [3] [5] [6] [7] [4]. Additionally, augmented late positive potential (LPP) responses have been observed, suggesting sustained attention to emotional stimuli [8] [9] [4].

However, not all studies report these findings, with some research indicating reduced ERP amplitudes in both early and late processing stages among anxious individuals. For example, Frenkel and Bar-Haim [10] found that early ERP components, such as the P1, P2, and early posterior negativity (EPN), did not significantly differ between anxious and non-anxious groups. Moreover, they observed reduced LPP amplitudes in response to fearful versus neutral stimuli in individuals with anxiety. Steinweg et al. [11] reported diminished discrimination of fearful faces during both early (EPN) and late (LPP) components, and Holmes et al. [12] noted enhanced early P1 response to fearful faces in individuals with high trait anxiety, followed by reduced amplitudes in subsequent ERP components. These mixed results may reflect variations in experimental paradigms, task demands, and analytical approaches [11].

Importantly, such inconsistencies may also point to underlying disruptions in attentional and emotional control mechanisms characteristic of anxiety. According to Attentional Control Theory [13], anxiety shifts the balance between the goal-directed (frontal-mediated) and stimulus-driven (occipital-mediated) attention, impairing top-down regulation of emotional responses.

Contemporary models further emphasize interactions among large-scale networks and predictive mechanisms in emotion perception [14, 15, 16]. For example, predictive coding frameworks propose that the brain continuously generates affective predictions that interact with sensory input. Del Popolo Cristaldi et al. showed that emotion regulation strategies modulate neural activity at multiple stages of this predictive processing hierarchy [14]. Other studies highlight the dynamic coupling between ‘core’ occipital regions and extended frontal-parietal systems during face processing, maximizing the efficiency of their communication [15], as well as the involvement of sensorimotor regions in embodied emotion recognition [16]. Collectively, these findings suggest that emotion perception arises from a combination of bottom-up visual encoding and top-down modulation by higher-order control networks. Disruptions to this balance may cause anxious individuals to show either sustained engagement with threatening stimuli— reflected in enhanced LPP due to reduced frontal disengagement—or, conversely, attenuated engagement indicative of avoidance strategies, as reflected in reduced LPP responses.

To resolve these discrepancies and clarify the underlying neural mechanisms, novel methodological approaches are required. Fast Periodic Visual Stimulation (FPVS) paradigms have recently emerged as a promising alternative to traditional ERP methods for investigating facial expression processing. FPVS leverages frequency-tagged stimuli to elicit steady-state visual evoked potentials (SSVEPs), providing an objective, high signal-to-noise ratio (SNR) measure of neural discrimination at pre-defined oddball frequencies [17] [18] [19] [20].

FPVS offers distinct advantages over traditional ERP techniques, including higher sensitivity, greater objectivity, and the ability to capture the complete neural response spectrum, encompassing both low-level sensory processing and higher-order cognitive functions [19] [17] [21] [22]. Notably, FPVS can probe implicit facial processing and has been shown to discriminate facial identity and emotion at a conceptual level, as evidenced by reduced or absent responses to inverted faces [23] [24]. Prior studies employing FPVS have demonstrated robust neural responses to emotional expressions such as fear, disgust, and happiness, with distinct activation patterns over occipital and occipito-temporal regions [24, 25]. Moreover, top-down attention has been shown to amplify early face responses; for instance, Baudouin et al. [26] found that explicit recognition of facial expressions enhanced early FPVS-EEG signals over occipital and frontal areas.

Despite these advances, FPVS has not yet been applied to investigate differences in facial expression processing between high-anxious and low-anxious individuals. The current study aims to fill this gap by examining neural reactivity to emotional facial expressions using the FPVS paradigm. This approach holds the potential to uncover novel insights into emotion processing deficits in anxiety disorders, contributing to a deeper understanding of their underlying neurobiological mechanisms. Furthermore, due to its robustness and efficiency, with interpretable data at the individual level and short recording times, FPVS shows promise as a practical tool for clinical assessment and screening [27].

Based on the literature, we hypothesized that high trait anxiety would be associated with larger neural discrimination responses to negatively-valenced faces. Specifically, we expected the high-anxiety group to show enhanced oddball responses at occipital electrodes for angry, fearful and sad facial oddballs than the low-anxiety group. No group differences were predicted for happy faces. Additionally, based on evidence of altered network interactions in anxiety, we aimed to explore potential disruptions in top-down connectivity. We therefore hypothesized that individuals with higher anxiety might show impaired anterior-to-posterior communication, reflected in reduced frontal-to-occipital response ratios.

## 2. Methods

### 2.1 Participants

Forty-one adults (ages 18-28, mean age = 20.8, 21 females) were recruited from the Mississippi State University student population. Ethical approval was obtained from the Mississippi State University Institutional Review Board, and all participants provided written informed consent. Participation was voluntary, and individuals could withdraw at any time. Data from three participants were excluded due to technical issues (two due to EEG recording failures and one due to excessive noise) resulting in a final sample of 38 participants.

Due to the exploratory nature of the current study, sample size was determined based on available time and economic resources, and no a priori power analysis was conducted. However, previous FPVS research informed our target of approximately 40 participants. For example, Poncet et al. [25] found robust FPVS responses to facial expressions with only 15 participants. Similarly, studies examining facial expression or identity processing in special populations have used sample sizes of 32 to 46 participants (e.g., [28] [29] [30]). Given this context, our final sample of 38 participants falls within the range of established FPVS paradigms, particularly for preliminary group comparisons.

Participants first completed a demographic questionnaire, which included self-reported anxiety diagnoses. They then completed the PROMIS Anxiety Short Form (Level 2) and the PROMIS Depression Short Form (Level 2). Participants were classified into the high-anxiety group if they either self-reported an anxiety diagnosis (n = 13) or had a PROMIS Anxiety t-score greater than 60 (n = 9 or an additional n = 4), resulting in a total of 17 individuals in the high-anxiety group (10 females; mean t-score = 60.7, sd = 7, range 43.3 – 72.9). The remaining 21 participants were classified as low-anxiety (11 females; mean t-score = 50.2, sd = 5, range 42.1 – 57.6). Only two participants scored above 60 on the PROMIS Depression scale, both of whom were in the high-anxiety group. All participants had normal or corrected-to-normal vision and hearing. In our statistical analyses, we examined both categorical group differences (high vs. low anxiety) and associations with anxiety as a continuous variable using the PROMIS t-scores.

### 2.2 Stimuli

Facial stimuli were selected from the validated FACES database [31], which includes 2,052 photographs of 171 individuals (85 female) aged 19 to 80, each displaying six basic facial expressions. For the present study, we selected two images per individual for five expressions (neutral, angry, fearful, sad, and happy) yielding a total of 1,710 unique images used. Disgust was excluded because it was not relevant to our hypotheses about anxiety and threat. We chose anger and fear as classic threat-related expressions, sadness as another negative emotion linked to dysphoria, and happiness as a positive contrast. Neutral faces served as the base stimuli. All images were resized from their original resolution (819 × 1,024 pixels) to 158 × 197 pixels (width × height; 96 dpi). On screen, each image subtended a visual angle of 5° × 6.25° (w × h) and was presented on a 24-inch monitor with a 60 Hz refresh rate using custom PsychoPy scripts [32].

### 2.3 Procedure

Participants were positioned 80 cm from the monitor. During the FPVS paradigm, facial stimuli were presented for 83 ms, followed by an 83 ms blank interval before the next image appeared. Stimuli were displayed in sequences of five images, with the first four being base images and every fifth image serving as an oddball stimulus. This design elicited two distinct steady-state responses: one at the base frequency (6 Hz) and another at the oddball frequency (1.2 Hz). Therefore, each of the four conditions (happy, sad, fearful, angry oddballs) was presented once as one continuous 122LJsecond FPVS sequence (∼732 faces per sequence; ∼146 oddballs). Conditions were shown in random order. Participants maintained fixation on a central cross and performed a color-change detection task (blue-to-red change for 400 ms, 6–8 times per run) to ensure engagement with the display without explicitly attending to facial expressions.

### 2.4 EEG recording

EEG data were recorded using a 64-channel BioSemi ActiveTwo system (BioSemi, Amsterdam, Netherlands) with a sampling rate of 256 Hz. The reference was set to the average of the left and right mastoids.

### 2.5 EEG processing

EEG preprocessing was performed using MATLAB (MathWorks Inc.) and the EEGLab toolbox. Data were cropped into segments including 2 seconds before and 3 seconds after each sequence, resulting in 127-second segments. A bandpass FIR filter (0.1-100 Hz) was applied. Channels exceeding five standard deviations from mean kurtosis values were identified and replaced via spherical interpolation. On average, fewer than 3% of channels were interpolated per participant (M=2.7, SD = 2.1). Data were re-referenced to the common average of all electrodes.

FPVS paradigms are relatively resistant to artifacts, facilitating the measurement of meaningful neural responses without the need for explicit correction of blinks or other artifacts [17]. Consequently, we chose not to apply artifact rejection methods such as Independent Component Analysis (ICA). This approach enhances the objectivity and reproducibility of the analysis and allows for easy automation, thereby making FPVS particularly suitable as a rapid screening tool.

The preprocessed data segments were trimmed to an integer number of 1.2 Hz cycles, excluding the first two cycles after sequence onset to avoid contamination from transient onset responses. A fast Fourier transform (FFT) was applied to extract normalized amplitude spectra across all channels and participants [24]. Prior research has demonstrated strong steady-state visual evoked potential (SSVEP) responses at oddball frequencies and their harmonics, with increased detection reliability when incorporating multiple harmonics [20] [33].

Significant harmonics were identified by computing z-scores relative to the amplitude mean and standard deviation of the 20 neighboring FFT bins (10 bins on each side, excluding immediately adjacent and extreme values) [24] [34]. A z-score threshold of 1.64 (p < 0.05, one-tailed) was applied to determine statistical significance [35] [36] [37]. Group-level analyses examined the number of significant harmonics across all electrodes and within predefined regions of interest (ROIs). Based on previous studies investigating FPVS brain responses to facial expressions (e.g. [24, 25]) ROIs included occipital (O: Oz, O1, O2), and lateral occipito-temporal (left lOT: PO7, P7, P9; right rOT: PO8, P8, P10). In order to investigate anterior contributions and top-down interactions we also included parieto-occipital (PO: POz, PO3, PO4), centro-parietal (CP: CPz, CP1, CP3) and frontal (F: Fz, FCs, AFz) ROIs. The range of significant harmonics for each condition was determined as the last harmonic meeting statistical significance. Harmonics corresponding to the base frequency (e.g., 6 Hz, 12 Hz) were excluded [38].

To quantify facial expression discrimination at oddball frequencies, baseline-corrected amplitudes (BCA) were computed by subtracting the mean amplitude of 20 surrounding bins (excluding immediately adjacent and extreme values) from the oddball frequencies and harmonics. Significant harmonics were identified up to 14 harmonics in pooled data and up to 20 harmonics in individual participants. As including non-significant harmonics should have minimal influence on summed BCA values, the BCAs were summed across the first 20 harmonics. Responses were quantified across all channels and predefined ROIs. Finally, the summed BCA spectrum was then scaled by dividing the value at each electrode by the mean across all electrodes.

Additionally, signal-to-noise ratio (SNR) was calculated for each frequency of interest as the amplitude value divided by the mean amplitude of the 20 neighboring bins (excluding adjacent and extreme values). SNR spectra were used for visualization, and grand averages of SNR and BCA were computed for each condition and electrode.

### 2.6 Statistical analysis

For the statistical analyses, we applied linear mixed models in JASP version 0.19.3 [39] with the summed BCA as the dependent variables. We fitted separate models for the four facial expressions: Angry, Fear, Happy, and Sad. ROI (F, CP, PO, O, lOT, rOT) and group (high-anxious, low-anxious) were added as fixed factors. We added depression as a covariate with minimal impact on the results. To examine anxiety as a continuous variable, we also rerun the linear mixed models with ROI and Anxiety t-score as the fixed factors. To account for the repeated testing, we included a random intercept per participant. Planned posthoc contrasts were corrected using Bonferroni. We decided to also look at ratios between top-down control regions (F and CP) and bottom-up perceptual regions (O, lOT, rOT) as indication of dysfunctional attentional and emotional control processes. For this we fitted separate linear mixed models for the following ratios: F/O, F/lOT, F/rOT, CP/O, CP/lOT, CP/rOT. The summed BCA ratios were the dependent variables with group and expression as the fixed factors. Additionally, we calculated Pearson’s r correlation coefficients between the BCA and Anxiety t-scores for the different expressions, ROIs and ratios.

## 3. Results

Analysis revealed strong responses at the oddball frequency (1.2 Hz) and its harmonics, as reflected in high signal-to-noise ratio (SNR) values across all conditions. Figure 2 visualizes clear oddball discrimination responses in the four facial expression conditions at the oddball frequency and its harmonics for the occipital ROI. All four facial expressions (angry, fearful, happy, and sad) exhibited significant mean z-scores above the threshold (z > 1.64) at oddball frequencies and their harmonics across all electrodes and within each ROIs. It can be seen that the high-anxious group typically had increased responses for all expressions compared to the low-anxious group. In spite of this, none of the linear mixed models showed a significant main effect of group. All linear mixed models did show significant main effects of ROI (p < 0.001 for all except p = 0.028 for sad). Considering the planned contrasts, however, both angry (p = 0.027) and sad (p = 0.023) facial expressions showed significantly increased responses for the high-anxious compared to low-anxious individuals at the occipital ROI. These differences remained statistically significant after controlling for depression as a covariate (p = 0.4 for angry and p = 0.29 for sad).

**Figure 1.**
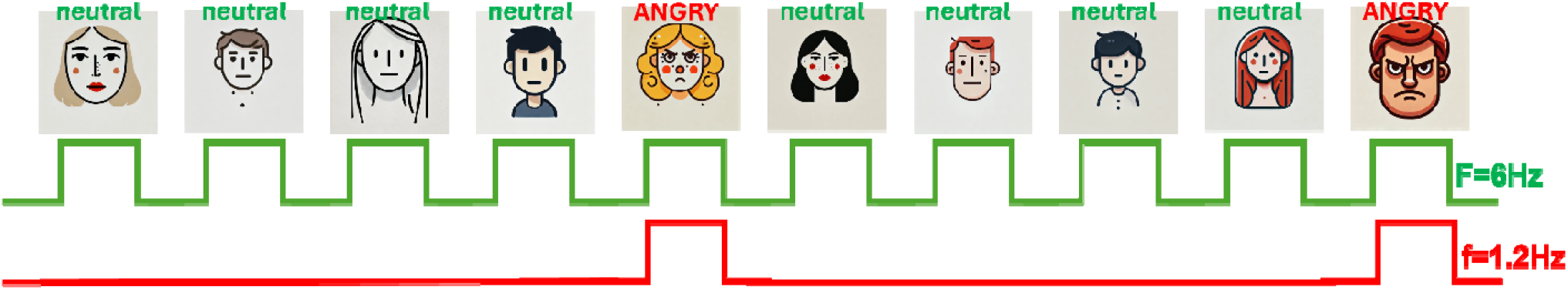
Schematic illustration of the experimental paradigm. Stimuli are presented at a rate of 6 Hz. In each 122 s stimulation sequence, base stimuli are selected from a large pool with oddball images presented at fixed intervals of one every fifth stimuli or 1.2 Hz. The example in this figure corresponds to condition D, angry faces. The faces shown here is just for illustration, we used real photos in the experiments.

**Figure 2.**
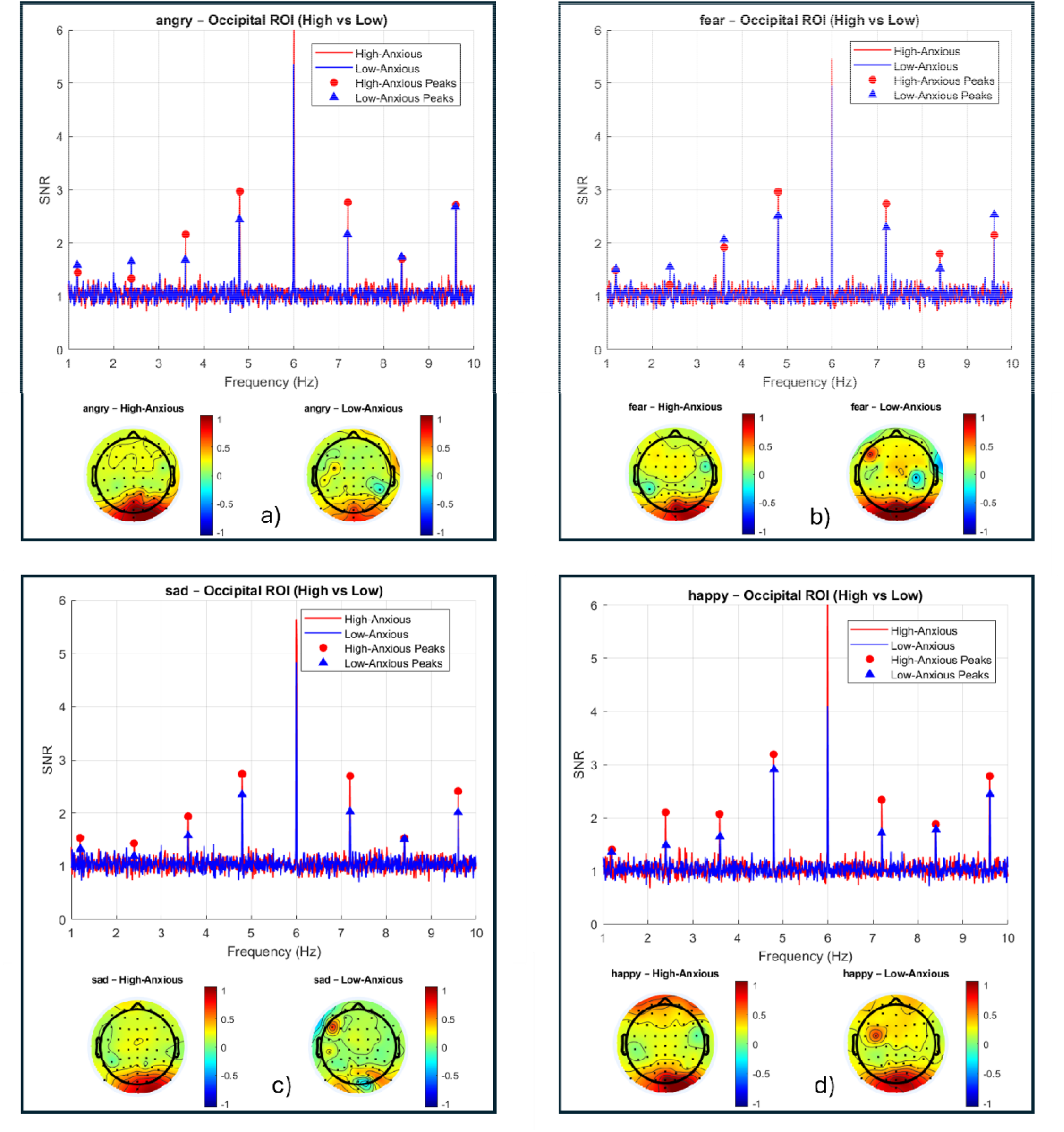
Oddball responses visualized via SNR spectra at the occipital ROI and BCA scalp maps for a) Angry facial expression, b) Fear facial expression, c) Sad facial expression, and d) Happy facial expression. Note that one outlier at CP1, determi ed by looking at z-scores > 3*sd, was removed for the scalp maps. Removing this participant had minimal impact on the statisti al findings.

Looking at the anterio-posterior ratios, a significant main effect of group was found for CP/rOT (F(1,35) = 4.673, p= 0.038, Cohen’s d ≈ 1.83) with the high-anxious individuals (M=0.386) showing significantly reduced ratios compared to the low-anxious individuals (M=0.521). Although not statistically significant, we also found a Pearson’s r of 0.155 for the CP/lOT ratio with a p = 0.064 indicating a possible trend.

## 4. Discussion

Our findings provide preliminary evidence of the potential promise of FPVS for assessing facial expression processing with ecologically valid and highly variable stimuli. This study demonstrated robust neural responses to emotional faces at both individual and group levels within short, two-minute recording sessions per condition, without requiring artifact removal in post-processing.

Importantly, we observed neural differences between high- and low-anxiety individuals in response to emotional faces. While preliminary, these findings suggest that FPVS may be sensitive to anxiety-related variation in facial emotion processing. Our classification of “high-anxiety” included individuals who either reported a prior anxiety diagnosis (some with t-score < 60) or scored above the clinical threshold (t-score > 60) on the PROMIS Anxiety scale. However, most participants exhibited relatively low current symptom levels, suggesting that the observed differences may reflect residual or trait-level vulnerability rather than acute anxiety. Further research with larger and more diverse samples is needed before any strong conclusions can be drawn about its broader applicability.

Theoretical models of anxiety often emphasize attentional biases, proposing that anxious individuals exhibit heightened vigilance toward negative and threat-related stimuli and difficulty disengaging from them [40] [41]. A consistent finding supporting the notion of hypervigilance is enhanced P1 components to threatening and sad facial expressions [3] [12] [4]. Our findings provide preliminary FPVS support for this notion as we saw enhanced BCA responses in occipital ROIs to angry and sad faces in high-anxious individuals. Given the implicit nature of the FPVS paradigm, these increased responses are likely to reflect early-stage pre-conscious processing [42]. This early enhancement may be driven by impaired feedback mechanisms from the amygdala modulating visual cortical activity [12].

However, in contrast to several previous studies (e.g. [10, 12, 43], we did not observe anxiety-related increases in neural responses to fearful oddballs. This finding aligns with Bar-Haim et al. [3], who reported that high-anxious participants showed increased P2 amplitudes only for angry faces, but not for neutral or other emotional expressions, including fear. Similarly, Steinweg et al. [11] found reduced discrimination between fearful and neutral faces in both early (EPN) and late (LPP) ERP components in individuals with high trait anxiety during a perceptual discrimination task.

One potential factor contributing to the discrepant findings is variation in methodological approaches, particularly the selection of facial expression stimuli. A plausible, though speculative, reason for the attenuated neural differentiation of fearful expressions in our study may stem from how fear was visually represented in the specific stimulus set employed. The FACES database used in this experiment did not require models to display surprise expressions, unlike other commonly used databases. As a result, some of the fear expressions may have unintentionally included visual elements typically associated with surprise (such as widened eyes or an open mouth). This overlap may have led to occasional misinterpretation of fear as surprise, potentially accounting for the observed differences. We note here that although the FACES images have been validated previously [31], we did not independently confirm recognition rates in our sample, which could be a confounding factor.

Another potential explanation for the absence of significant anxiety-related effects for fearful faces involves the nature of fear as a social signal. Fearful expressions often indicate the presence of an ambiguous or unspecified threat in the environment, rather than a direct one. This ambiguity may elicit hypervigilance toward all surrounding stimuli, including neutral faces, thereby reducing the neural differentiation between oddball and base stimuli in the fear condition [44, 42]. In this scenario, anxious individuals may allocate increased attention to both fearful and neutral faces, leading to a diminished contrast between them and thus a reduced oddball response. In contrast, angry faces typically signal a direct interpersonal threat, particularly when gaze is directed at the observer, thereby eliciting more selective attention toward the angry (oddball) stimuli specifically [44]. This could explain why heightened responses in high-anxious individuals were observed for angry but not fearful expressions in our study.

As emotional processing extends into later stages, individuals with higher anxiety often show divergent patterns of neural activity. Some studies report elevated P3/LPP amplitudes in response to negative or threatening faces, indicative of sustained emotional engagement [4] [45]. Others observe reduced neural reactivity in these same components, interpreted as reflecting avoidance strategies that suppress later-stage processing [11] [12] [10]. Our study did not identify significant differences between high- and low-anxious groups in occipito-temporal or fronto-central ROIs across emotional expressions. Several explanations are plausible: high-anxious individuals may have adopted a hypervigilance-then-avoidance strategy, thereby suppressing later neural responses; alternatively, they may have perceived all faces, including neutral ones, as threatening, resulting in reduced differential responses [46]. Another possibility is that the relatively brief 833 ms interval between oddball stimuli allowed for overlap of successive LPPs, obscuring clear identification of these late-stage components.

Regardless of the precise mechanism, the divergence in LPP findings across studies may reflect two facets of the same dysregulated emotional system. Enhanced LPPs can signify persistent engagement due to impaired frontal inhibition, while attenuated LPPs may reflect dysfunctional avoidance strategies, both underpinned by deficient top-down control. Consistent with this interpretation, our finding of reduced central-parietal to right occipito-temporal ratios in the high-anxious participants suggests disrupted functional integration between early perceptual regions and higher-order regulatory areas. This pattern may reflect impaired cortico-limbic regulation, including dysregulated interactions among the amygdala, prefrontal cortex, and associated structures implicated in emotional control and stress response [47].

While this study provides preliminary evidence supporting the feasibility of using FPVS to assess facial expression processing in clinical populations, several limitations should be acknowledged. First, although our sample size is comparable to other FPVS studies involving special populations, it was limited in both size and diversity. Participants were primarily college students, which may constrain generalizability. However, given the high prevalence of psychopathology, including suicidality, among college students across multiple universities, this population remains relevant for investigating depression and anxiety [48]. Nevertheless, future studies should aim to recruit larger, more heterogeneous samples, particularly including individuals actively seeking mental health care.

A second limitation is that anxiety diagnoses were based on self-reported prior diagnosis and PROMIS Anxiety t-scores, rather than structured clinical interviews. Incorporating clinician-administered assessments would enhance diagnostic accuracy and allow for clearer delineation of comorbidities.

Third, we only observed significant group differences using categorical classifications, whereas analyses treating anxiety as a continuous variable yielded no statistically significant associations. One possible explanation is that many participants in the high-anxiety group, while reporting a prior diagnosis, showed low current symptom severity, possibly due to remission or treatment. This discrepancy may reflect the limited variability in anxiety scores across our sample, with most participants falling in a moderate range and few extreme scores. It is well known that categorical groupings can inflate effect sizes [49], and as such, these findings should be interpreted with caution. They serve only as tentative, exploratory evidence for the utility of FPVS in detecting anxiety-related neural differences. Future research should apply this paradigm to samples with broader and more clinically diverse symptom profiles to support dimensional analyses and improve generalizability.

Fourth, while our study broadly addressed anxiety, we did not assess subtypes (e.g., social anxiety, generalized anxiety, panic disorder). Different subtypes may exhibit distinct neural profiles, and future work should explore whether FPVS can differentiate among these. Moreover, expanding FPVS applications to other conditions such as PTSD or depression could clarify whether emotion-processing deficits are shared or disorder-specific.

Finally, a methodological limitation involves the spatial resolution of EEG. Our findings are based on scalp-recorded responses and interpreted in terms of functional topographies (e.g., occipital as perceptual, frontal as control-related). However, EEG cannot precisely localize neural sources, particularly in deeper or overlapping regions. Future studies employing source reconstruction techniques or combining FPVS with neuroimaging modalities such as MEG or fMRI may provide clearer insight into the anatomical generators underlying emotion-related neural responses.

The FPVS approach has previously been used to investigate facial and emotional sensitivity in children with autism spectrum disorder [29] [28]. Our findings provide initial evidence that FPVS may be able to differentiate anxious from non-anxious individuals based on their neural responses. Given its high signal-to-noise ratio, rapid acquisition, and interpretability at the individual level, FPVS is a promising approach for basic emotion research and may have future utility in clinical screening. All raw EEG data and analysis scripts from this study have been made publicly available on the Open Science Framework (OSF; https://osf.io/5xkae/files/osfstorage) to facilitate transparency and replication.

